# Accelerating *k*-mer-based sequence filtering

**DOI:** 10.1101/2025.06.16.659853

**Authors:** Igor Martayan, Léa Vandamme, Bede Constantinides, Bastien Cazaux, Charles Paperman, Antoine Limasset

## Abstract

**Motivation:** The exponential growth of global sequencing data repositories presents both analytical challenges and opportunities. While *k*-mer-based indexing has improved scalability over traditional alignment for identifying relevant documents, pinpointing the exact sequences matching numerous queries remains a hurdle. In particular, searching for numerous *k*-mers with a single large query or multiple distinct queries strains existing exact matching tools, whose performance scales poorly with an increasing number of patterns. At the same time, indexing entire vast datasets for infrequent or ad-hoc searches is often resource-prohibitive. Designing fast methods for matching a large number of *k*-mers without exhaustive pre-indexing is therefore critical.

**Contributions:** We propose an efficient solution to the problem of *k-mer-based sequence filtering*: given a set of *k*-mers of interests and a threshold, quickly evaluate whether an arbitrary sequence has a number of *k*-mer matches above or below the threshold. Our approach demonstrates how minimizer-based based sketching, alongside SIMD acceleration, can enhance the performance of streaming searches, and is implemented as a Rust tool named *K2Rmini*. On a consumer laptop, K2Rmini is able to filter long reads at 2 Gbp/s.

**Availability:** https://github.com/Malfoy/K2Rmini.

## 1 Introduction

The immense volume of available sequencing data presents both significant opportunities and challenges in bioinformatics. Assembled datasets have reached the Petabase scale, as seen in the 2 Petabytes of unitigs from the Logan project [6], while even curated databases like GenBank are nearing 6 Terabases [35]. Although holistic analyses of these large-scale datasets, such as those on viral [11], soil [24], gut [29], and environmental metagenomes [30] are promising, they remain computationally expensive. Identifying matches to a query sequence in large collections, a fundamental task exemplified by BLAST [4], has evolved. Traditional alignment tools were overwhelmed by data growth, leading to the use of strings of fixed length *k*, or *k*-mers [43]. To improve scalability, indexing structures allowing false positives were adopted, making shared *k*-mer fingerprints a necessary, though not sufficient, condition for a match [26,27,22]. Some methods further enhance scalability by indexing only a fraction of *k*-mers, relying on detecting substantial matches from partial signals [10,33,1].

Large-scale indexes can thus efficiently identify a small, relevant subset of documents (e.g., read sets, unitigs, contigs). However, this initial filtering isn’t final. Due to false positives from probabilistic indexes, or sparse/non-collinear matches, queried *k*-mers may not be truly present or usable for mapping. Therefore, recovering specific sequences within identified documents that genuinely share *k*-mers with the query is crucial. This refinement, moving from relevant documents to relevant sequences suitable for alignment, mitigates downstream scalability issues and can reduce data volume by orders of magnitude, such as identifying a few relevant sequences from billions.

The problem of *k-mer-based sequence filtering* can be formalized as follows: given a set of *k*-mers of interest 𝒬 = {*q*_1_, …, *q*_*m*_} (the “queries”), a set of sequences 𝒮 = {*S*_1_, …, *S*_*n*_} *T* and a threshold, output for each *S*_*i*_ ∈ 𝒮 whether ∑_*q*∈*𝒬*_ #occ(*q, S*_*i*_) ≥ *T*, i.e. whether *S*_*i*_ contains at least *T* occurrences of *k*-mers of interest. In practice, the threshold can either be set as an absolute value *T* ∈ ℕ or as a relative value proportional to the size of the sequence: given *t* ∈ [0, 1], *T*_*i*_ = ⌈*t* · (|*S*_*i*_| − *k* + 1)⌉.

### 1.1 Related work

Although sequence-level indexes have been proposed [20,40], indexing and storing entire dataset collection at this resolution remains expensive and challenging at scale, thus favoring lightweight filtering methods directly from unindexed data. Efficient multi-pattern matching is a highly studied topic from both the theoretical [8] and practical point of view [13]. Practical implementation can be found in classical data processing with grep-like efficient tools such as Ripgrep [14] or Hyperscan [41]. Some tools perform pattern matching adapted to genomic data, such as Seqkit [37,38], fqgrep [19] or grepq [9]. However, these solutions tend to scale poorly as the number of query *k*-mers increases and fail to leverage the fixed length of the search patterns. This is a critical issue, as querying numerous *k*-mers is often necessary for large queries (contigs, long reads, whole genomes), searching for many distinct sequences (variant or gene collections), or optimizing batched queries. The recent BackToSequences [5] tool, used in the Logan project, addresses these issues by simply indexing query *k*-mers in a hash table to search for target documents. Very recently, both Deacon [7] and Cleanifier [42] were proposed as scalable solutions for contaminant depletion, that is removing sequences originating from a given genome, which is a direct application of sequence filtering. In particular, Deacon effectively uses many of the techniques presented in this article. Finally, while to our knowledge it has not been used for sequence filtering, the spectral Burrows-Wheeler transform (SBWT) [3] is also a suitable method for this problem thanks to efficient matching statistics [25].

### 1.2 Contributions

This work presents three main contributions. First, we introduce a new algorithm that uses random minimizers to quickly filter out sequences that contains too few *k*-mers of interest, effectively reducing the cost of negative matches by a factor *w/*2. Second, we propose an optimized implementation named *K2Rmini*, that takes advantage of vectorized instructions to parse sequences, hash *k*-mers and compute minimizers efficiently. Third, we compare our approach to many different tools and evaluate their scalability for a large number of patterns, highlighting the benefits of different methods. Our tool, K2Rmini, is able to filter 2 Gbp/s on a consumer laptop and is available online at https://github.com/Malfoy/K2Rmini.

## 2 Methods

Classical multi-pattern matching algorithms, which generalize the Knuth-Morris-Pratt approach, construct deterministic finite automata to achieve linear time complexity relative to the pattern and input data sizes. While these methods offer strong theoretical guarantees, they often exhibit poor cache locality. Furthermore, they address the more general problem of matching variable-length patterns, whereas our work focuses exclusively on fixed-size patterns. This constraint creates an opportunity for hash-based techniques, which are better suited for large pattern sets and more amenable to hardware acceleration through vectorized instructions.

One of the state-of-the-art solutions, BackToSequences [5], uses an efficient hash table containing all *k*-mers of interest and skips irrelevant fields. However, its main limitation is its necessity to perform a hash table lookup for every *k*-mer in the data stream. This process becomes computationally expensive, especially when the hash table is large and does not fit into low-level caches. To address this bottleneck, we propose K2Rmini (https://github.com/Malfoy/K2Rmini), a method that integrates minimizer-based filtering and Single Instruction, Multiple Data (SIMD) operations.

### 2.1 Using minimizers to upper bound the number of *k*-mer matches

Minimizers [36,34] are a widely used *k*-mer sampling method that selects the smallest *m*-mer out of every *w* consecutive one, usually using a random ordering. The set of sampled minimizers is typically much smaller, by a factor 2*/*(*w* + 1) called the *density* [32], than the whole sequence while preserving a lot of information, making it a key component of large-scale sequence analysis. An important property of minimizers is that they are *locally consistent*: two sequences sharing contexts of *w m*-mers will always sample the same minimizers from these contexts. Thus, comparing minimizers can be used as a proxy to identify similar sequences [23,10,12].

One core idea of our approach is to leverage this property to avoid exhaustive *k*-mer scanning. Given a set 𝒬 of *k*-mers of interest, we compute the set of associated minimizers ℳ(𝒬) = {minimizer(*q*) : *q* ∈ 𝒬}. Then, instead of checking every *k*-mer from a sequence *S*, we only compare the minimizers of *S* against ℳ(𝒬). We know that each minimizer match implies that up to *w k*-mers of *S* are in 𝒬. Thus, if we have *ℓ* minimizer matches, the number of *k*-mer matches is upper-bounded by *ℓ* × *w*, so having 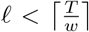 is sufficient to discard a sequence.

While this upper bound is quite accurate for sparse matches, it is very loose (by a factor close to 2) for dense matches. One way to refine this upper bound is to observe that if two minimizers are *d* positions apart from each other, they cover at most *w* + *d k*-mers. Therefore, if we have a chain of consecutive minimizer matches at positions *p*_1_, …, *p*_*r*_, the number of *k*-mer matches is upper-bounded by *w* + *p*_*r*_ − *p*_1_.

In practice, we often have access the exact number of *k*-mers covered by each minimizer as a byproduct of the sliding window algorithm [17]. In that case, we can simply sum them to get a tight upper bound.

### 2.2 Minimizer-based filtering algorithm

Our algorithm, detailed in Algorithm 1, works in two passes. The first pass computes an upper bound on the number of *k*-mer matches, as detailed in the previous section, by performing 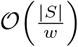 queries to 𝒯_ℳ_ (the hash table of minimizers). This has two main advantages: not only do we reduce the number of lookups, but querying 𝒯_ℳ_ is also faster than 𝒯_𝒬_ (the hash table of *k*-mers) because it is roughly *w/*2 times smaller when *k*-mers from 𝒬 are consecutive.

The second pass only happens for sequences with an upper bound on *k*-mer matches above the threshold. In that case, we must count the exact number of *k*-mer matches to identify true hits. Deacon [7], another tool tailored for decontamination developed by some of the authors, uses a simplified version of the method proposed here. In particular, it omits this second pass, reducing computational overhead at the cost of potential false positives.

#### Algorithm 1

Minimizer-based filtering

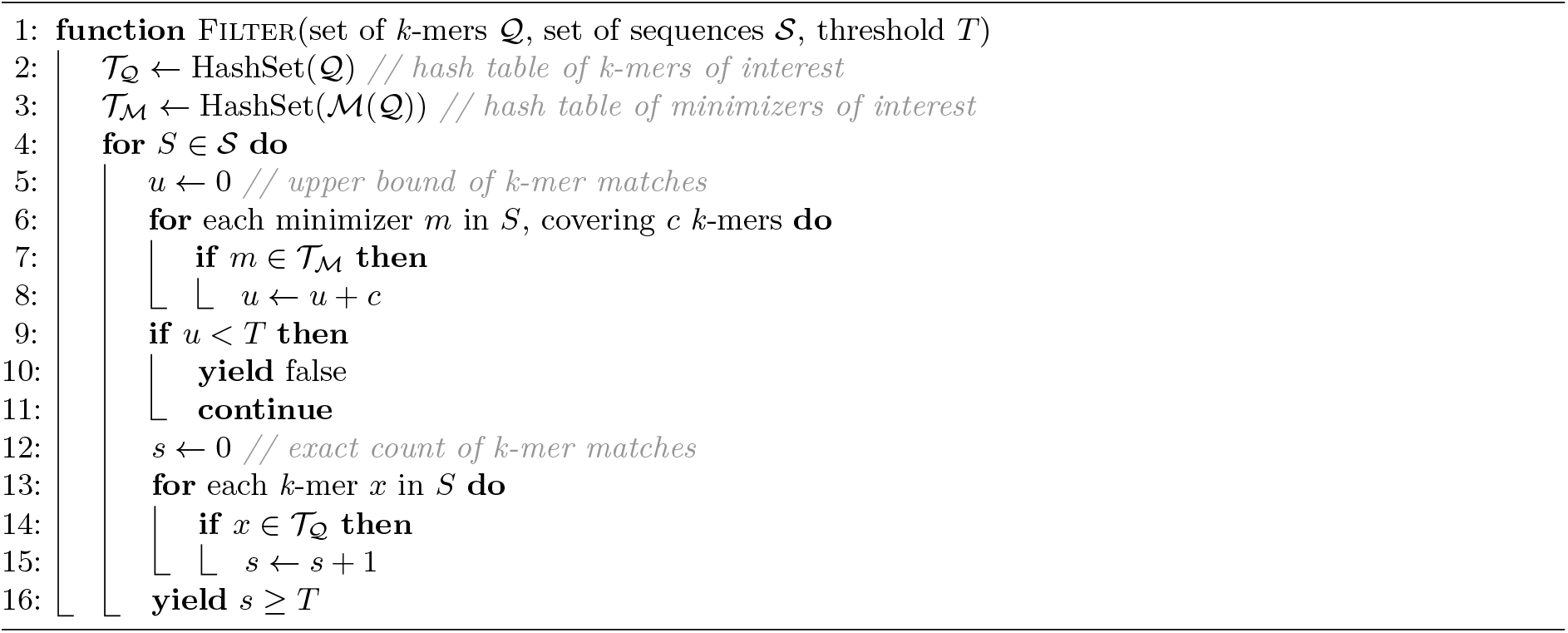

### 2.3 Vectorization

Multiple parts of our algorithm are accelerated using vectorized instructions. First of all, we implemented a library called `helicase` (https://github.com/imartayan/helicase) that vectorizes the parsing of the sequence files and outputs bitpacked representations for faster processing. Then, we build upon SimdMinimizers [17] to have a vectorized computation of minimizer positions, along with the number of *k*-mers that they cover. SimdMinimizers relies on a completely branchless sliding window minimum algorithm using two stacks and processes 8 independent chunks of the sequence within a single SIMD register. Finally, we use a vectorized rolling hash adapted from NtHash [28] to accelerate the *k*-mer lookups in the second pass.

### 2.4 Parallelization

On top of the vectorized computation of the matches, we adopt a parallelization strategy for the main loop based on a producer-consumer model. A producer thread parses the sequences and sends batches of sequences (up to a desired length) to multiple consumer threads, which in turn compute the matches on their batch and output the results. One approach that we have not implemented yet but could be worth trying would be to let each consumer thread directly parse a chunk of the input file in parallel, thus removing the need for a producer thread and reducing the communication overhead.

## 3 Results

All experiments were run with the benchmarking workflow available at https://github.com/imartayan/K2Rmini_experiments. The benchmarking framework is easily extensible, since each additional program only requires a dedicated command wrapper to be added to a common interface before it can be executed and plotted with the other tools. All reported times include both the cost of indexing the queried patterns and the filtering step. Benchmarks were performed on a dual-socket Intel Xeon Gold 6430 machine with 64 cores and 64 GB of RAM running Ubuntu 24.04.2. Because no ready-to-use SBWT-based sequence filtering implementation was available for our setting, we implemented the SBWT baseline ourselves^4^ using the Rust sbwt crate (https://docs.rs/sbwt/latest/sbwt/).

### 3.1 State-of-the-art analysis

To evaluate state-of-the-art tools, we benchmarked a selection of specialized and general-purpose programs while varying the number of queried *k*-mers. The specialized tools included Seqkit 2.13.0 [37,38], fqgrep 1.1.1 [19], grepq 1.6.5 [9], BackToSequences 0.8.3 [5], Cleanifier 1.2.0 [42], Deacon 0.14.0 [7], and our SBWT-based implementation. For comparison, we also included grep 3.11, ripgrep 15.1.0 [14], and Hyperscan 5.4.2 [41] (or Vectorscan 5.4.12 on ARM). As shown in Figures 1 and 2, we benchmarked these tools on 2.6 Gbp of PacBio HiFi reads from HG002^5^ while increasing the number of queried *k*-mers.

**Fig. 1:**
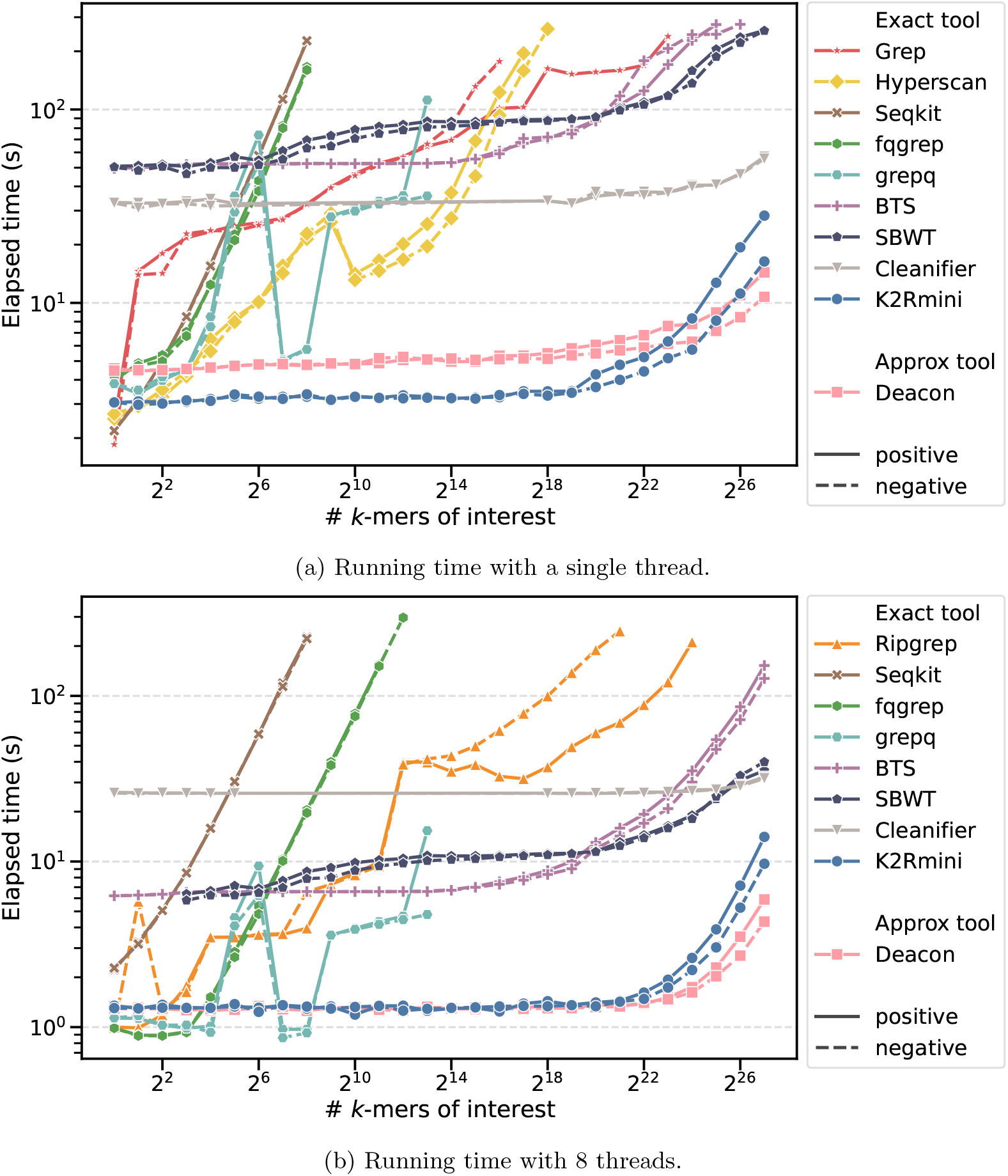
Running time of state-of-the-art tools when filtering 2.6 Gbp of PacBio HiFi reads from HG002 (`m54328_180928_230446`) with an increasing number of queried *k*-mers for *k* = 31. Positive *k*-mers are extracted from the reads while negative *k*-mers are random sequences.

**Fig. 2:**
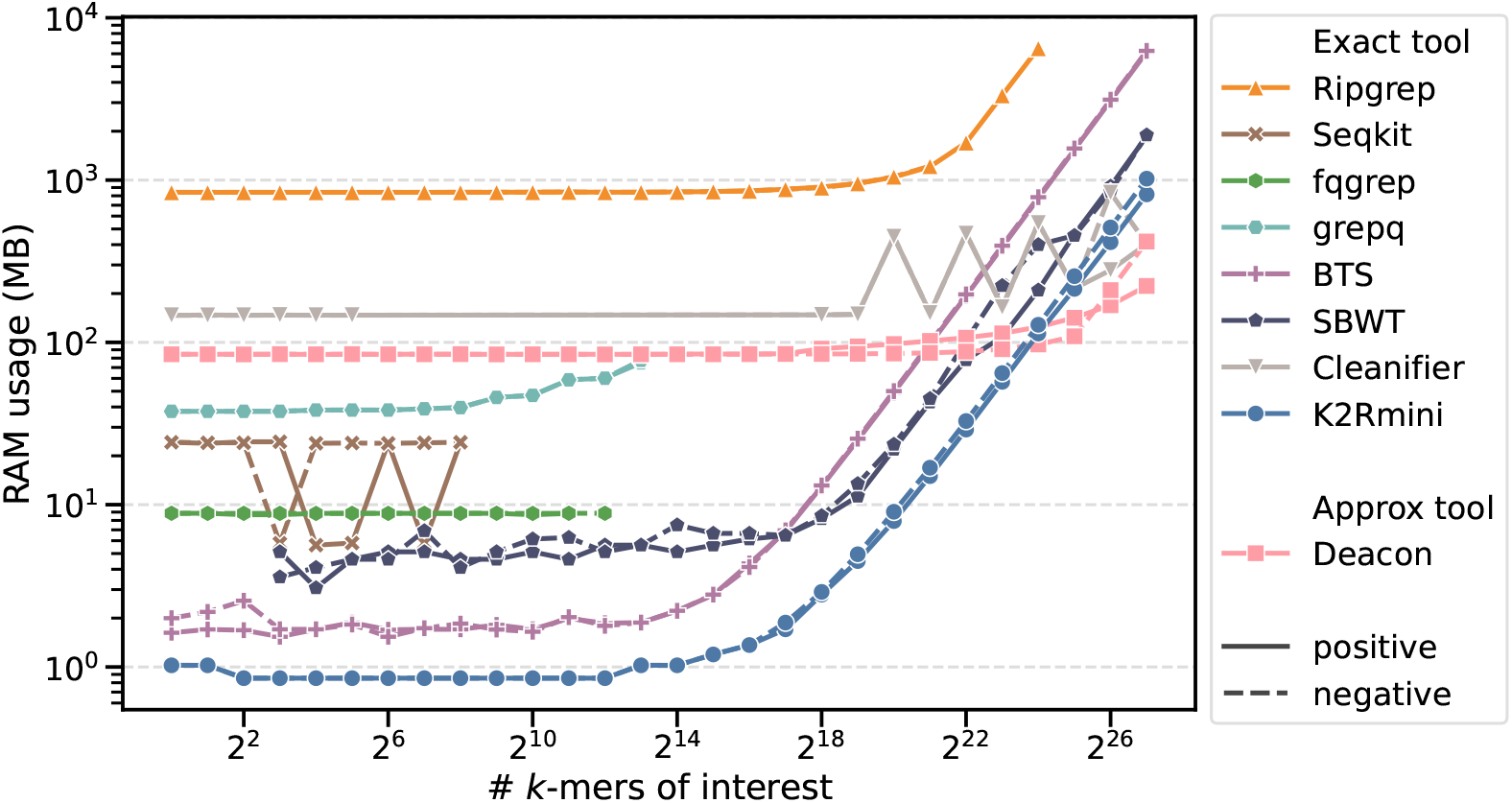
Memory usage of state-of-the-art tools when filtering 2.6 Gbp of PacBio HiFi reads from HG002 (m54328_180928_230446) with an increasing number of queried *k*-mers for *k* = 31. Positive *k*-mers are extracted from the reads while negative *k*-mers are random sequences.

The results separate the tested methods into two broad groups. Classical multiple-pattern matching tools such as grep, ripgrep, Hyperscan, Seqkit, fqgrep, and grepq scale poorly as the number of queried *k*-mers increases. Their running time rises rapidly and they become impractical once the query set reaches the large regimes that matter for genomic filtering. In contrast, methods that explicitly index the queried patterns, or a sketch derived from them, remain effective over the full range of tested query sizes.

Among these scalable approaches, Deacon and K2Rmini are the fastest overall. Cleanifier is also largely insensitive to the number of queried *k*-mers, but with a clearly higher absolute running time. SBWT and BackToSequences remain much more scalable than general-purpose pattern matchers, although both become slower than Deacon and K2Rmini as the number of queried *k*-mers grows. In the multithreaded setting, SBWT remains consistently ahead of BackToSequences for the largest query sets, while both are clearly outperformed by the minimizer-based approaches.

The gap between positive and negative queries is especially informative for K2Rmini. For negative *k*-mers, most reads are rejected during the minimizer pass, which keeps the running time close to flat across a broad range of query counts. For positive *k*-mers, more reads pass this first filter and trigger the exact counting phase, so the running time increases more substantially. This behavior directly reflects the design of K2Rmini, where minimizers are used to avoid unnecessary exact searches, but exact verification is still required whenever the upper bound is above threshold.

This comparison must be interpreted more carefully for Deacon. Deacon only indexes minimizers and does not perform an exact second pass on the surviving reads. Its excellent scalability therefore comes from solving a relaxed version of the problem, which may introduce false positives. By contrast, K2Rmini uses minimizers as a lossless prefilter and performs exact *k*-mer counting whenever necessary.

A practical observation is that at some point, pattern indexing itself is a visible part of the total running time and filtering is no longer the only bottleneck.

Memory usage reveals a similar separation between approaches. General-purpose tools such as ripgrep already have a large baseline memory footprint and can reach several gigabytes for the largest query sets. By contrast, the specialized genomic tools initially use much less memory, but their scaling behavior differs substantially. Among the scalable exact methods, K2Rmini has the lowest memory footprint over most of the explored range and remains below BackToSequences and SBWT at large query sizes. BackToSequences starts from a very small footprint on small query sets, but its memory usage grows steeply and eventually becomes the highest among the indexed genomic methods. SBWT scales more smoothly than BackToSequences, but still requires substantially more memory than K2Rmini for large pattern sets. Cleanifier and Deacon exhibit a higher baseline memory usage, yet their growth is more moderate, which is consistent with their good scalability in running time. Overall, these results show that K2Rmini provides the best end-to-end compromise among exact methods, combining low running time with low memory usage over a broad range of query sizes.

### 3.2 Influence of parameters

To better understand the behavior of the most scalable approaches, we compared BackToSequences, SBWT, Deacon, and K2Rmini while varying the number of threads and the *k*-mer size. The results are shown in Figures 3 and 4.

**Fig. 3:**
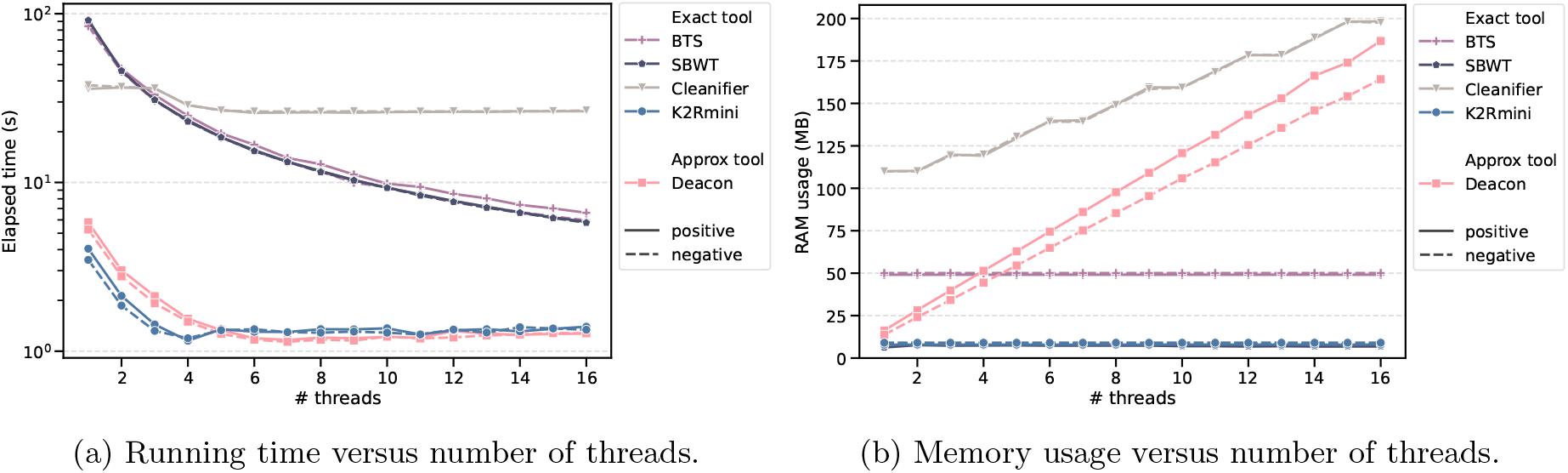
Influence of the number of threads when filtering 2.6 Gbp of reads with 1M patterns of size *k* = 31.

**Fig. 4:**
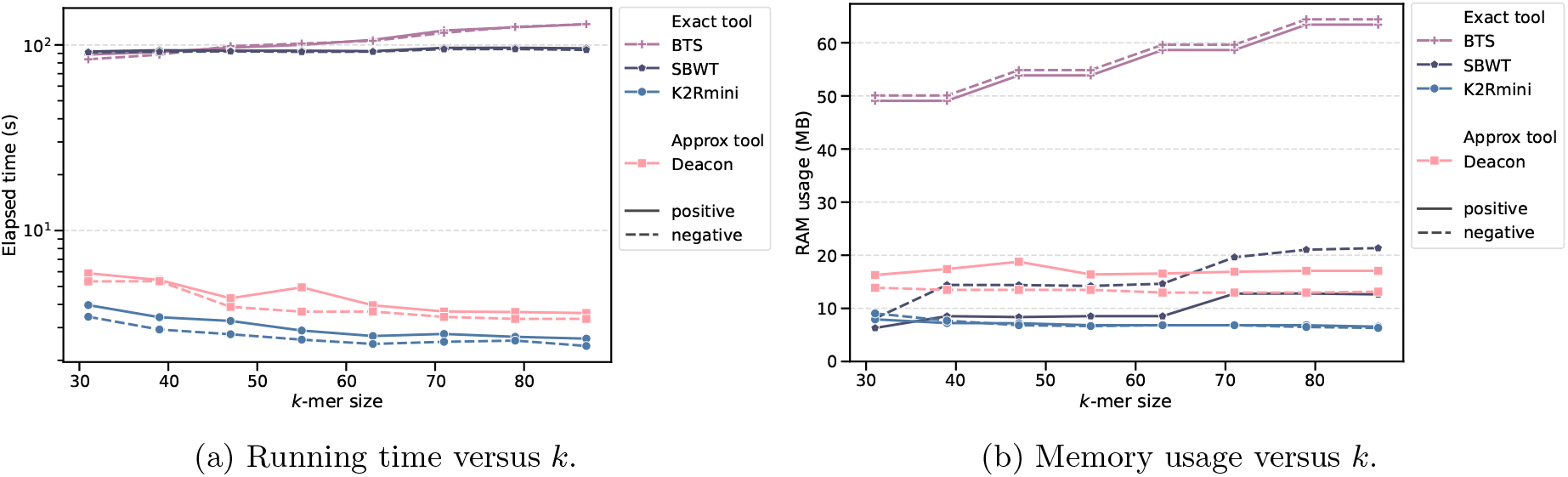
Influence of the *k*-mer size when filtering 2.6 Gbp of reads with 1M patterns and a single thread. Cleanifier is not included because it does not support *k >* 32.

When increasing the number of threads, K2Rmini remains the fastest method throughout the benchmark. Most of the speedup is obtained within the first four threads, after which the running time stabilizes around 1.3–1.4 seconds. Deacon shows a very similar trend, with slightly higher running times and the same early saturation. This suggests that, for both minimizer-based approaches, the main bottleneck quickly shifts away from the matching procedure itself toward parsing and other shared overheads. In contrast, BackToSequences and SBWT continue to benefit from additional threads over the full range, with their running times decreasing steadily from about 100 seconds on one thread to around 6 seconds on 16 threads. Cleanifier shows almost no parallel speedup in this experiment, remaining close to 35 seconds regardless of the thread count.

The memory trends differ markedly from the time trends. K2Rmini has the lowest memory footprint and remains nearly constant as the number of threads increases, staying around 8–10 MB. SBWT also remains essentially flat, at about 8 MB. BackToSequences has a larger but still stable footprint, around 50 MB, independent of the number of threads. By contrast, Deacon and Cleanifier both show a substantial increase in memory usage with additional threads. Deacon grows roughly linearly from about 15 MB on one thread to around 180 MB on 16 threads, while Cleanifier increases from about 110 MB to about 195 MB. Therefore, among the tested methods, K2Rmini combines the best running time with the smallest and most stable memory usage in the multithreaded setting.

We also evaluated the effect of the *k*-mer size with a single thread and 1M queried patterns. As *k* increases, K2Rmini becomes slightly faster, decreasing from about 4 seconds at *k* = 31 to about 2.3 seconds for the largest tested values. This trend is consistent with our minimizer-based design. Keeping the minimizer size fixed while increasing *k* enlarges the minimizer window, reducing minimizer density, and therefore decreasing the number of lookups required during the first filtering pass. Deacon exhibits a similar, though less pronounced, decrease in running time. In contrast, BackToSequences becomes slower as *k* increases, rising from about 80 seconds to more than 120 seconds, while SBWT remains roughly constant around 90 seconds. These results again highlight that minimizer-based filtering benefits from larger *k*-mer sizes, whereas exact indexed lookup of every *k*-mer does not.

The memory usage as a function of *k* remains moderate for all methods, but the trends are again distinct. K2Rmini stays almost constant around 7–8 MB across the full range of tested values. Deacon remains below 20 MB, with only a small increase as *k* grows. SBWT increases more noticeably, specifically after each power of two (32 and 64 in the plot), from about 7–8 MB at small *k* to about 20 MB for the largest tested values, especially on negative queries. BackToSequences has the highest memory usage throughout this experiment, steadily increasing from about 50 MB to more than 60 MB. Overall, these experiments confirm that K2Rmini is not only the fastest of the exact methods considered here, but also the most memory-efficient and the least sensitive to both thread count and *k*-mer size.

### 3.3 Real data

To complement the synthetic benchmarks, we compared BackToSequences and K2Rmini on several real datasets with heterogeneous sequence lengths and structures, summarized in Table 1. The evaluation includes Oxford Nanopore ultra-long reads (SRR23365080), PacBio HiFi reads of 10–20 kb (SRX7897685– 8), and Illumina 250 bp reads (ERR3239454). We also evaluated assembled sequences from the Logan project [6], including both contigs and unitigs (SRR7853572), and the complete human T2T reference genome (GCF_009914755.1), which represents very long contiguous sequences. These experiments were run on a laptop equipped with a 13th Gen Intel Core i9-13950HX processor, 64 GB of RAM, and an SSD, using 8 threads. For the positive case, the queried patterns were sequences of length 1000 selected from each dataset, using 100 patterns for short reads and unitigs so as to total 10^6^ queried bases. For the negative case, the queried patterns were random sequences of length 1000. Across all datasets and in both positive and negative settings, K2Rmini is consistently faster than BackToSequences in both CPU time and wall-clock time.

**Table 1:** Runtime comparison between K2Rmini and BackToSequences (BTS) using 8 threads.

| Experiment | CPU time (s) |  |  | Elapsed time (s) |  |  |
| --- | --- | --- | --- | --- | --- | --- |
|  | K2Rmini | BTS | Speedup | K2Rmini | BTS | Speedup |
| ONT positive | 1850.65 | 19529.04 | ×10.55 | 531.83 | 2626.34 | ×4.94 |
| ONT negative | 959.72 | 17021.16 | ×17.74 | 125.13 | 2013.64 | ×16.09 |
| HiFi positive | 279.46 | 7527.53 | ×26.94 | 33.11 | 907.56 | ×27.41 |
| HiFi negative | 273.75 | 4716.76 | ×17.23 | 33.70 | 557.88 | ×16.55 |
| Illumina positive | 278.24 | 1537.61 | ×5.53 | 43.56 | 173.02 | ×3.97 |
| Illumina negative | 267.35 | 1597.82 | ×5.98 | 40.41 | 180.89 | ×4.48 |
| T2T positive | 55.85 | 223.67 | ×4.00 | 8.88 | 37.19 | ×4.19 |
| T2T negative | 9.81 | 148.98 | ×15.19 | 2.31 | 25.63 | ×11.10 |
| Contigs positive | 14.05 | 102.10 | ×7.27 | 2.18 | 12.66 | ×5.81 |
| Contigs negative | 13.83 | 102.29 | ×7.40 | 2.15 | 12.70 | ×5.91 |
| Unitigs positive | 76.56 | 191.45 | ×2.50 | 13.87 | 26.49 | ×1.91 |
| Unitigs negative | 75.29 | 192.07 | ×2.55 | 14.07 | 26.65 | ×1.89 |

The largest gains are observed on ONT and HiFi reads. On ONT data, K2Rmini is 10.55× faster than BackToSequences in CPU time and 4.94× faster in elapsed time for positive queries, while the speedup increases to 17.74× and 16.09× on negative queries. On HiFi reads, the gains are even larger for positive queries, reaching 26.94× in CPU time and 27.41× in elapsed time, and remain very high on negative queries with speedups of 17.23× and 16.55×. On Illumina reads, the improvement is more moderate but still substantial, ranging from 5.53× to 5.98× in CPU time and from 3.97× to 4.48× in elapsed time. Overall, these results confirm that the minimizer-based prefilter is especially effective on raw reads, with the strongest gains on long-read datasets.

The same trend holds on assembled and reference sequences, although with more variable gains. On the T2T genome, K2Rmini is 4.00× faster in CPU time and 4.19× faster in elapsed time for positive queries, while negative queries lead to much larger speedups of 15.19× and 11.10×. On Logan contigs, the gains are stable between positive and negative cases, around 7.3× to 7.4× in CPU time and 5.8× to 5.9× in elapsed time. Logan unitigs show the smallest difference, with speedups around 2.5× in CPU time and 1.9× in elapsed time in both settings. More generally, negative queries tend to benefit more from K2Rmini than positive ones, which is consistent with the minimizer filter rejecting most sequences early and therefore avoiding the exact counting phase. Overall, K2Rmini consistently outperforms BackToSequences across all tested datasets, with the strongest improvements on long reads and on negative queries.

## 4 Discussion

This work has demonstrated that a search strategy based on SIMD-accelerated minimizer filtering is fast and resource-efficient across a broad spectrum of applications. Similar tools are being developed for screening sequence repositories for antimicrobial resistance mutations, emerging pathogen surveillance, or sequencing contaminant filtration [7]. Currently, the main limitation of our implementation is that the indexation of the patterns is single-threaded and relies on a generic hash table. One promising direction to improve the indexation step would be to implement a concurrent hash table that relies on minimizers for grouping keys, such as the one recently described in kache-hash [21]. This would be especially relevant since we already have efficient methods to compute both minimizers and *k*-mer hashes, and we could thus easily batch insertions and queries. A second direction for improvement would be to parallelize the parsing step by attributing independent chunks of the input file to each thread, which would fully benefit from our vectorized parsing library and reduce the communication overhead.

Future work could explore alternative search strategies, including using denser minimizer schemes to further reduce the number of minimizers to query [18,16,15,32,2]. We could also imagine a middleground between the costly exact search used in the second pass and doing no search at all at the cost of false positives, possibly by using small filters for faster queries with a lower false positive rate.

Finally, as the throughput of the core algorithm increases, performance bottlenecks shift to other components of the pipeline, mostly I/O bounded. While uncompressed FASTA/Q files are unnecessarily wasteful, reading input from compressed FASTA/Q files is a major bottleneck, highlighting the need for more performant alternatives [39,31].

## Acknowledgments

This work is funded by the French National Research Agency AGATE ANR-21-CE45-0012 and SxC ANR-24-CE25-2874-01. BC is supported by Wellcome Trust Award 313694/Z/24/Z.

## Footnotes

4 https://github.com/imartayan/K2Rmini_experiments/tree/main/sbwt_filter

5 https://s3-us-west-2.amazonaws.com/human-pangenomics/NHGRI_UCSC_panel/HG002/hpp_HG002_NA24385_son_v1/PacBio_HiFi/15kb/m54328_180928_230446.Q20.fastq

